# Demographic and immune-based selection shifts before and after European contact inferred from 50 ancient and modern exomes from the Northwest Coast of North America

**DOI:** 10.1101/051078

**Authors:** John Lindo, Emilia Huerta-Sánchez, Shigeki Nakagome, Morten Rasmussen, Barbara Petzelt, Joycellyn Mitchell, Jerome S Cybulski, Eske Willerslev, Michael DeGiorgio, Ripan S Malhi

## Abstract

The susceptibility of Native Americans to infectious disease has been postulated as a major factor for their population decline after European contact. To investigate if a preexisting genetic component contributed to this phenomenon, we analyzed 50 exomes of both ancient and modern individuals from the Northwest Coast of North America, dating from before and after European contact. We confirmed the genetic continuity between the ancient and modern individuals and modeled the population collapse after European contact, inferring a 57% reduction in effective population size. We also identified signatures of positive selection on immune-related genes in the ancient but not the modern group. The strongest selection signal in the ancients came from the human leukocyte antigen (HLA) gene *HLA-DQA1*, with alleles that are close to fixation. The important immune function of *HLA-DQA1* supports an ancient adaptation to the environments of the Americas. The modern individuals show a marked decrease in the frequency of the associated alleles (the most pronounced variant showing a 64% difference). This decrease is likely due to the environmental change associated with European colonization, which resulted in a shift of selection pressures, whereby negative selection may have acted on the same gene after contact. Furthermore, the selection pressure shift could correlate to the European-borne epidemics of the 1800s, suffered in the Northwest Coast region. This is among the first studies to examine a single population through time and exemplifies the power of such studies in uncovering nuanced demographic and adaptive histories.

---

The decline of Native American populations after European contact has been linked to several factors including warfare, alterations in social structure, and an overwhelming introduction of European-borne pathogens^1–3^. Although the extent of the population decline remains contentious, European-borne epidemics may have disproportionately contributed to the phenomenon^4,5^. The debate has prompted researchers to explore the possibility of genetic susceptibility, where low genetic variation in HLA genes and immunologically naïve populations are linked to the exacerbated pathogen-associated mortality rates^6–8^. Assumptions of homogeneity among certain immune genes^8^, however, are based on surveys of living Native Americans who represent the surviving members of communities affected by European contact and colonization. Thus, they fail to consider immune-related genetic factors that may have existed before contact.

The immunological history of the indigenous peoples of the Americas is undoubtedly complex. As people entered the Americas and expanded into different regions, approximately 15,000 to 20,000 years before present (BP)^9,10^, groups encountered environments with varying ecologies and with relatively little gene flow from other continental populations until European contact^11^. We hypothesize that indigenous peoples adapted to local pathogens, resulting in long-lasting changes to immune-related loci. Ancient immune adaptations are suspected to have occurred throughout human history as populations spread into varying environments across the globe^12^. If the indigenous peoples of the Americas adapted to local pathogens, those adaptations would have proven useful in ancient times but not necessarily after European colonialists altered the environment with their pathogens, some of which may have been novel^13–15^. Existing genetic variation as a result of adaptation prior to European contact could thus have contributed to the indigenous population decline after European contact.

## Results

### Samples and sequencing

To investigate possible immune-related genes under selection before European contact, we sequenced the exomes of ancient and modern First Nations individuals of the Prince Rupert Harbour (PRH) region of British Columbia, Canada (Supplementary Fig. 1). We then performed genomic scans for positive selection and functional characterization of genes exhibiting the strongest signals. Exomes of 25 modern individuals from two Coast Tsimshian communities, Metlakatla and Lax Kw’alaams (henceforth referred to as “Tsimshian”), were sequenced to a mean depth of 9.66x. The exomes of 25 ancient individuals from archaeological sites in the PRH region (henceforth referred to as “PRH Ancients”; Supplementary Fig. 1) were sequenced to a mean depth of 7.97x. Contamination estimates using the exome-wide data revealed a mean contamination of 0.94% with a 95% CI: 0.83-1.10 ^16^ (Supplementary Table 3). All 25 ancient individuals exhibited patterns of C→T and G→A transitions consistent with deamination due to post-mortem DNA damage^17,18^ (Supplementary Fig. 3). Mitochondrial haplogroups were determined for each ancient individual, all showing haplotypes previously identified in Native Americans^19^ (Supplementary Table 1).

### Examining the genetic relationship between the ancient and modern individuals

Before proceeding with selection scans we investigated the genetic relationship among the ancient and modern individuals to confirm continuity between the two groups. For these analyses, C/T and G/A polymorphic sites were removed to guard against biases resulting from DNA damage. Multidimensional scaling was performed to assess the genetic relationships of our samples to individuals from the 1,000 Genomes Project^20^, other Native American populations^21^, and two ancient individuals from the Americas^22,23^ (Fig. 1b). The analysis revealed an affinity among the PRH Ancients and the Tsimshian, with the Tsimshian drifting toward Europeans as expected from presumed European admixture^24^. We next employed *ADMIXTURE*^25^ to separate our samples and other worldwide populations into clusters. This analysis (at *K*=5 clusters) suggests that the Tsimshian are a mixture of ancestral components stemming from the PRH Ancients and Europeans (Fig. 1a). Next, the evolutionary relationship among our samples and other worldwide human populations was evaluated via *TreeMix*^26^. With a single migration event, the PRH Ancients exhibit minimal drift and appear ancestral to the Tsimshian with European admixture occurring between the two groups (Fig. 1c). These analyses, combined with local oral histories and evidence from archeology and mitogenomes^19^, allowed for the inference that the ancient and modern individuals represent a single population through time, which includes pre-and post-European contact.

**Figure 1.**
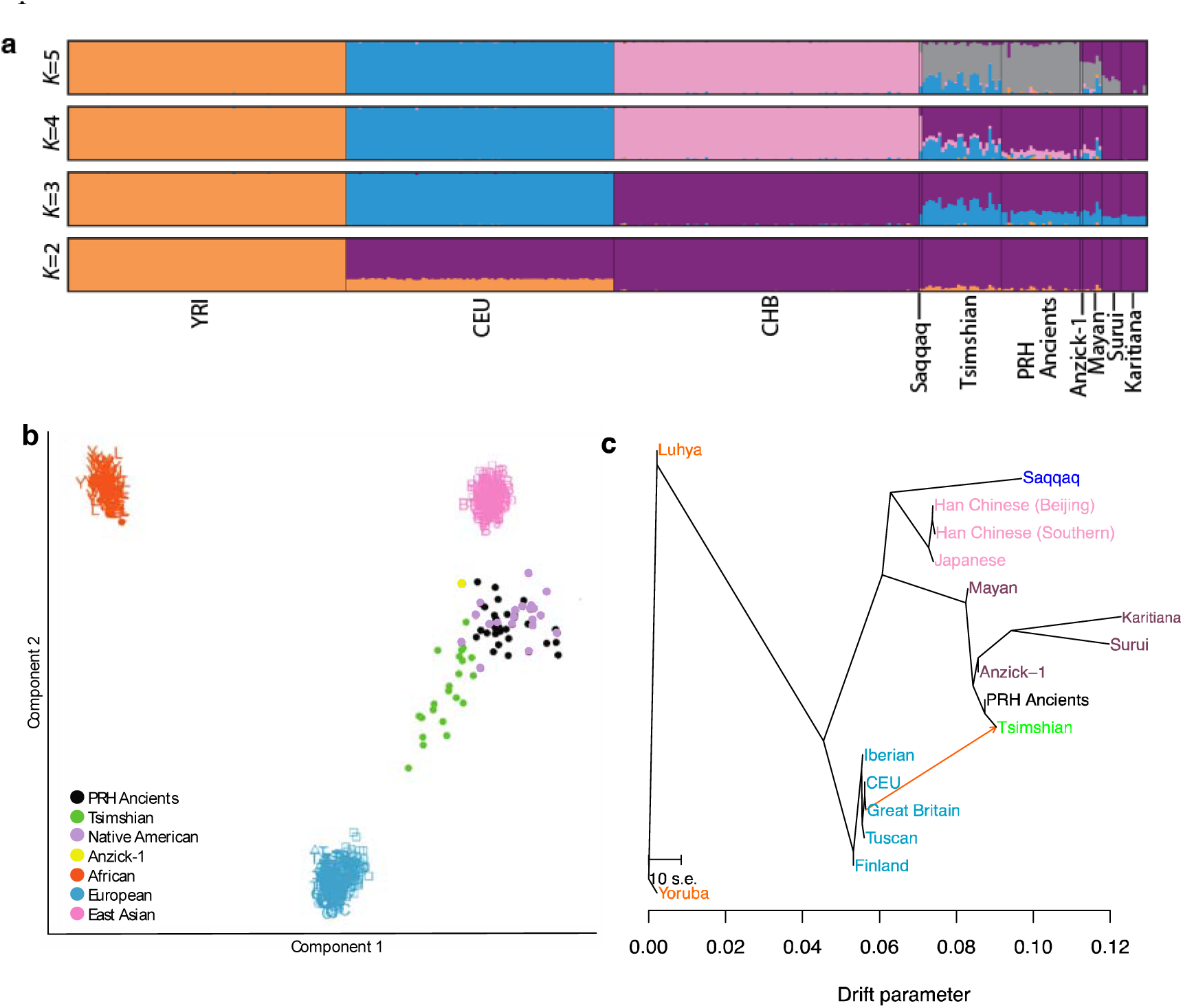
Population structure and demographic model of the PRH Ancients and Tsimshian. (a) *ADMIXTURE* analysis depicting ancestry proportions assuming the number of genetic components, *K*, is 2 to 5. The analyses included reference populations from the 1,000 Genomes Project Phase 2 dataset^20^, Native American populations sampled from the Karitiana, Surui, and Maya^21^, and two ancient samples from the Americas—the Saqqaq^22^ and Anzick-1^23^. (b) Multi-dimensional scaling plot. The Native American (Surui, Mayan, and Karitiana) populations fall with the PRH Ancients and the Anzick-1 ancient sample from Montana. The modern Tsimshian fall along the gradient leading from the Native Americans and the Europeans, reflecting their admixed history with Europeans. (c) *TreeMix* graph with a single admixture event.

### Demographic model

We also inferred the population history of the Tsimshian by taking into account the bottleneck that occurred after European contact^1,27,28^ (Fig. 2, see Methods). Utilizing both ancient and modern exome-wide data, the demographic parameters were inferred utilizing the joint derived site frequency spectrum of potential synonymous sites with respect to human reference genome hg19. The best-fitting model suggests that a bottleneck occurred approximately 175 years BP (bootstrap 95% confidence interval: 125-225, Table 1) in the ancestors of the modern Tsimshian with an accompanying reduction in effective population size of 57%. The timing of the bottleneck coincides with the documented smallpox epidemics of the 19^th^ Century and historical reports of large scale population declines^29,30^. A majority of the European admixture in the population likely occurred after the epidemics^24,29^.

**Figure 2.**
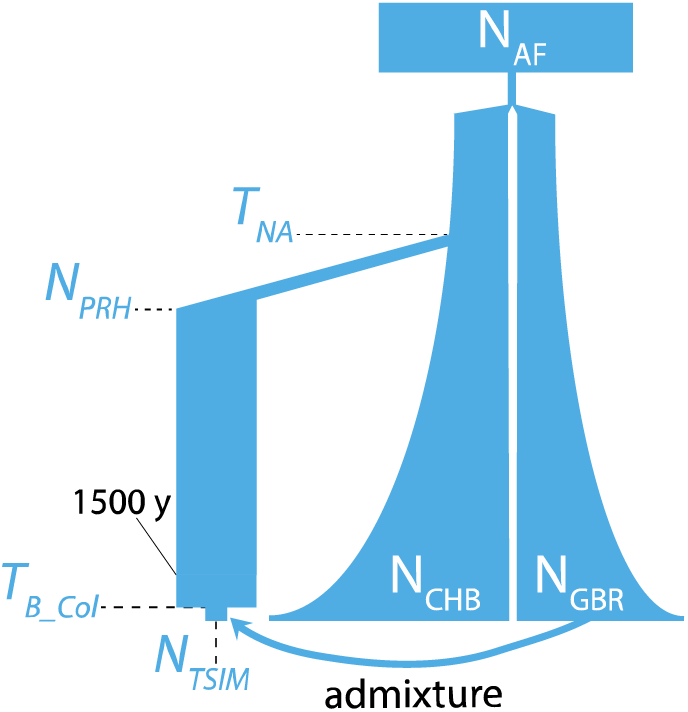
Demographic model. Best fitting model depicting the Tsimshian bottleneck after European contact a the subsequent admixture with Europeans. Fixed demographic parameters are from Gravel et al. ^31^ and admixtu parameters are from Verdu et al. ^24^. The parameters in blue were inferred with FastSimCoal2^32^. Population sizes and time splits are not shown to scale.

**Table 1.**
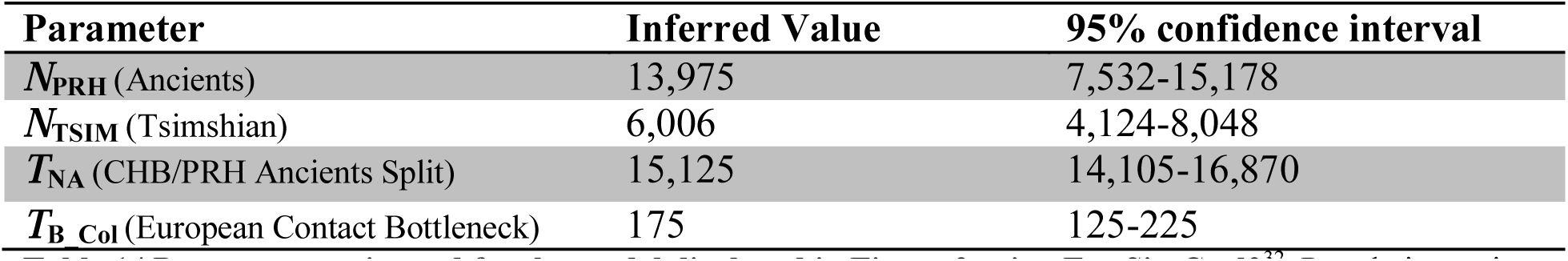
Parameters estimated for the model displayed in Figure 2 using FastSimCoal2^32^. Population estimates are for diploid individuals. Time estimates are in years, assuming a generation time of 25 years.

### Scan for positive selection

To safeguard against false-positive signals of positive selection due to the apparent admixture of Tsimshian individuals with Europeans, we performed an admixture correction (see Methods). The populations were scanned for selection signals, with and without correcting for admixture, utilizing the population branch statistic (PBS)^33^. PBS has proven effective in detecting positively-selected loci amongst high-altitude populations^33,34^. Twenty-five Han individuals from Beijing (CHB), part of the 1,000 Genomes Project^20^, served as the third comparative population. The statistic computes the amount of differentiation at a given locus along a branch leading to a specific population by comparing transformed *F*_ST_ values between each pair of three populations. Figure 3a displays population-specific differentiation for the mean across the exome and for our top candidate gene (*HLA-DQA1*) discussed below.

**Figure 3.**
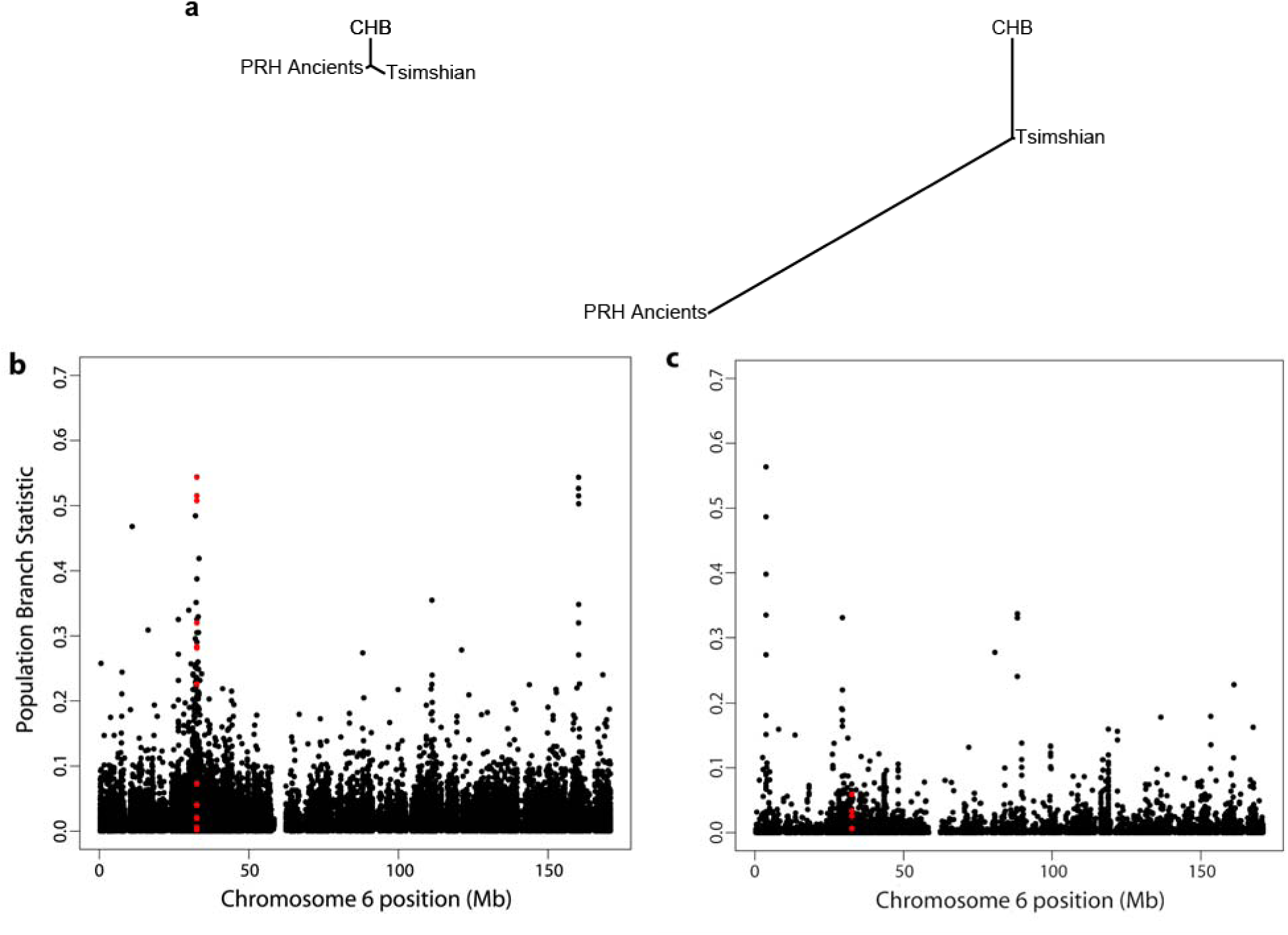
Population branch statistic (PBS) of the *HLA-DQA1* gene. (a) Trees based on exome-wide data, with the left tree showing a small branch length of the PRH Ancients relative to Tsimshian and CHB and the right tree showing strong differentiation along the PRH Ancients branch at the *HLA-DQA1* gene. (b,c) Manhattan plots of the PBS score as a function of SNP position on chromosome 6 for the PRH Ancients (b) and the modern Tsimshian (c). The region highlighted in red shows PBS scores for SNPs within 10kb of the *HLA-DQA1* gene.

The genes showing the most extreme and significant PBS values in the PRH Ancients represent strong candidates for positive selection, of which the top candidate, *HLA-DQA1*, is directly involved with immune function (Table 2). Enriched gene ontologies were also identified from the ranked list of genes generated from the PBS scan, which highlight immune function related to antigen presentation (Table 3). To assess whether the selection signals were extreme relative to expectations under neutrality, the PBS scores were compared to the distribution of scores based on neutral simulations using our inferred demographic history (Supplementary Fig. 11). Variants from the top candidate with the most pronounced frequency changes were confirmed via Sanger sequencing in all ancient samples reporting data (Supplementary Fig. 10).

**Table 2.** Genes with the strongest frequency changes in the PRH Ancient individuals.

| Gene | PBS | P-value | SNPs | Description |
| --- | --- | --- | --- | --- |
| *HLA-DQA1* | 0.275500845 | 0.01675 | 17 | Major histocompatibility complex, class II, DQ alpha 1 |
| *CELA3A* | 0.194966437 | 0.03102 | 29 | Basic leucine zipper nuclear factor 1 |

**Table 3.**
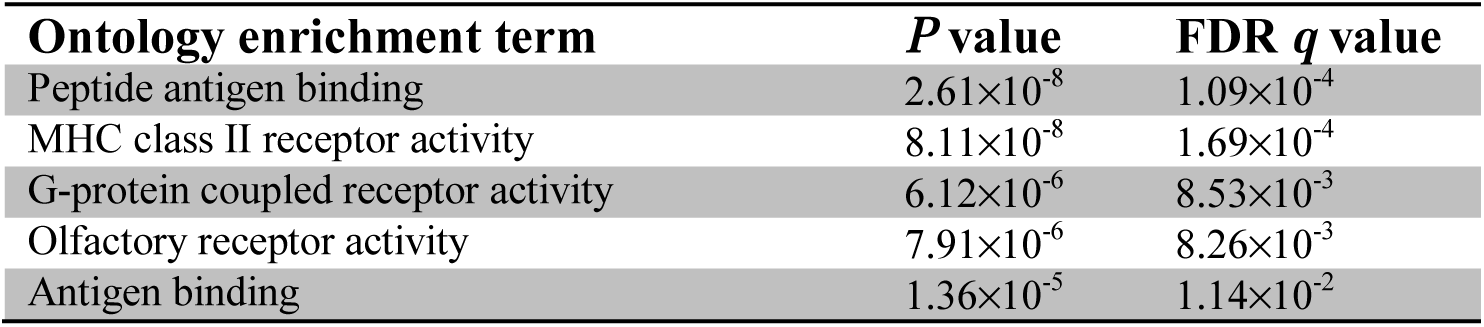
Enriched gene ontologies for the PRH Ancient individuals derived from the PBS selection scan ranked list. Calculated via GOrilla^35^.

| Ontology enrichment term | P value | FDR q value |
| --- | --- | --- |
| Peptide antigen binding | $2.61 \times 10^{-8}$ | $1.09 \times 10^{-4}$ |
| MHC class II receptor activity | $8.11 \times 10^{-8}$ | $1.69 \times 10^{-4}$ |
| G-protein coupled receptor activity | $6.12 \times 10^{-6}$ | $8.53 \times 10^{-3}$ |
| Olfactory receptor activity | $7.91 \times 10^{-6}$ | $8.26 \times 10^{-3}$ |
| Antigen binding | $1.36 \times 10^{-5}$ | $1.14 \times 10^{-2}$ |

### Relevance of the *HLA-DQA1* gene

The most extreme PBS score belonged to the *HLA-DQA1* gene, which encodes for the alpha chain of the major histocompatibility complex (MHC), class II, DQ1 isoform. The *HLA-DQA1* SNP with the most pronounced frequency difference between the PRH Ancients (100%) and the Tsimshian (36%) falls in the 5’ untranslated region (Table 4). This region may be indicative of selection acting on the regulation of the gene, as the associated alleles exhibit evidence of chromatin alterations and eQTL hits in a variety of cells—including Monocytes-CD14+ and Primary T helper 17 cells^36^ (Supplementary Table 10). The chromosomal region where the gene is located also shows strong differentiation along the branch leading to the PRH Ancients (Fig. 3).

**Table 4.**
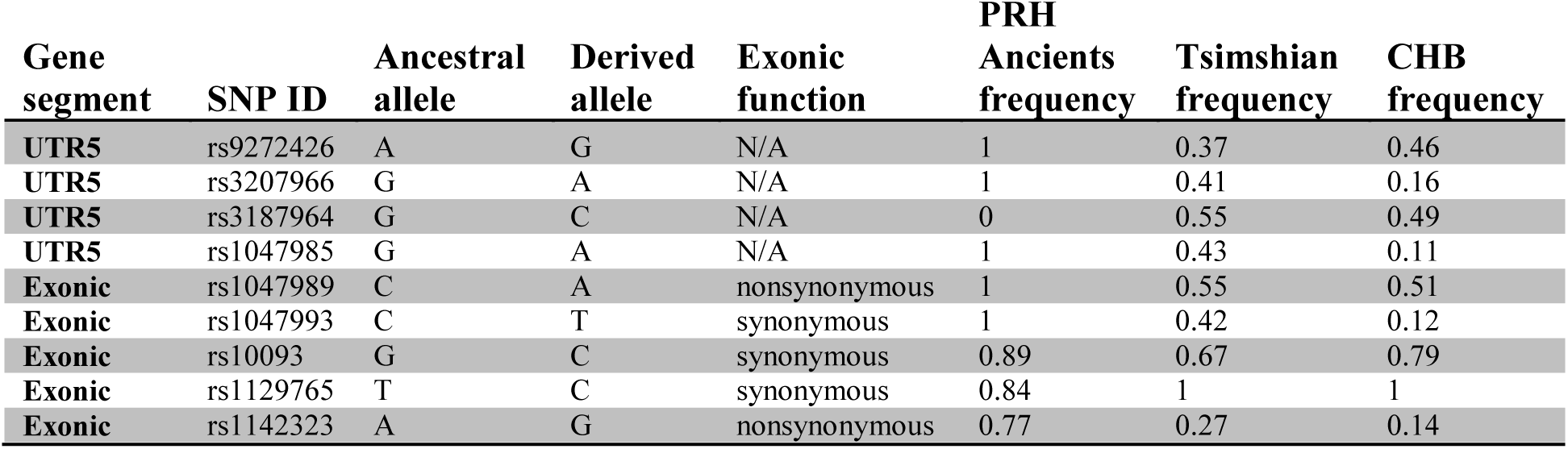
Population frequencies for the *HLA-DQA1* SNPs on chromosome 6. The population frequencies are for the derived allele with respect to the chimpanzee reference.

HLA-DQ is one of the three main types of MHC class II molecules, along with DR and DP, and is mainly expressed on antigen presenting cells^37^. MHC class II molecules are responsible for binding to extracellular pathogen peptides and presenting them to CD4+T helper cells, which activate a targeted adaptive immune response towards the associated microbe^38^. The molecules are known to be highly polymorphic, mainly due to sequence differences corresponding to the binding domain of the molecule, which can impact binding affinities^39^. Because of this variety in binding domains, differing MHC class II isoforms can have differing disease outcomes due to the restriction imposed on T-cell activation^38^. The polymorphic nature of these molecules across *different* populations, however, would not explain the heightened differentiation in the PRH Ancients with respect to their presumed descendants, the Tsimshian.

### Haplotype structure and local ancestry of *HLA-DQA1*

The top candidate for selection in the ancient population, *HLA-DQA1*, showed large allele frequency changes in the UTR5 region of the modern population (Table 4). Although there is a slight reference bias in the ancient samples due to mapping and possibly the design of our capture probes (Supplementary Fig. 14), the high frequency cannot be attributed to this feature since the derived alleles putatively under selection are for the *alternate* allele. To assess whether the frequency change was due to European admixture, we examined the haplotype structure among populations. To visualize the haplotypes in the *HLA-DQA1* region, we phased the ancient and modern samples using Beagle 4.1^40^. We took a randomly chosen haplotype from ancient sample PRH 125, and computed the number of pairwise differences to this haplotype for each haplotype in the modern and ancient samples as well as the Great Britain (GBR) samples from the 1,000 Genomes Phase 3 data^41^. We then ordered the haplotypes based on their number of pairwise differences to this arbitrarily chosen haplotype from sample PRH 125, and grouped them by population. Supplementary Figure 15a shows very similar haplotypes between the ancient and modern individuals, while those of the European population are distinct.

We next explored the local ancestry of the *HLA-DQA1* gene. We used RFMix^42^ to infer ancestry along chromosome 6 in the modern Tsimshian population. We utilized the PopPhased program, which tries to correct for phasing errors, and a window size of 0.2cM, four generations since the admixture event between the Tsimshian and Europeans, and 100 trees generated per random forest. For the reference panel, we used Phase 3 data from the 1,000 Genomes Project^41^. We used 25 individuals from the GBR (European panel), CHB (East Asian panel), and PEL (Native American Panel). The PEL chosen showed little to no admixture (see Methods). Supplementary Figure 15b indicates that only one haplotype could be attributed to European ancestry, while the remaining 49 are attributed to Native American ancestry.

### Simulations of the *HLA-DQA1* allele trajectories

To explore whether the allele frequencies differences between the two time periods could be explained by long-term balancing selection, drift, or changes in selection pressures, we conducted a series of simulations based on our demographic model. First, we examined whether long-term balancing selection under heterozygote advantage could explain our data. The parameters inferred from our demographic model were implemented in the forward-time simulator SLiM^43^. A *de novo* mutation was introduced 5 million years in the past (assuming a generation time of 25 years) that evolved under heterozygote advantage (per-generation selection coefficient *s*=0.1, and dominance parameter *h*=100) until the present. The distribution of the resulting PBS scores can be seen contrasted with the observed data in Figure 4a. Because the distribution of the PBS scores under long-term balancing selection is shifted toward small values compared to neutrality (Supplemental Figure 11), the data are inconsistent with long-term balancing selection under heterozygote advantage.

**Figure 4.**
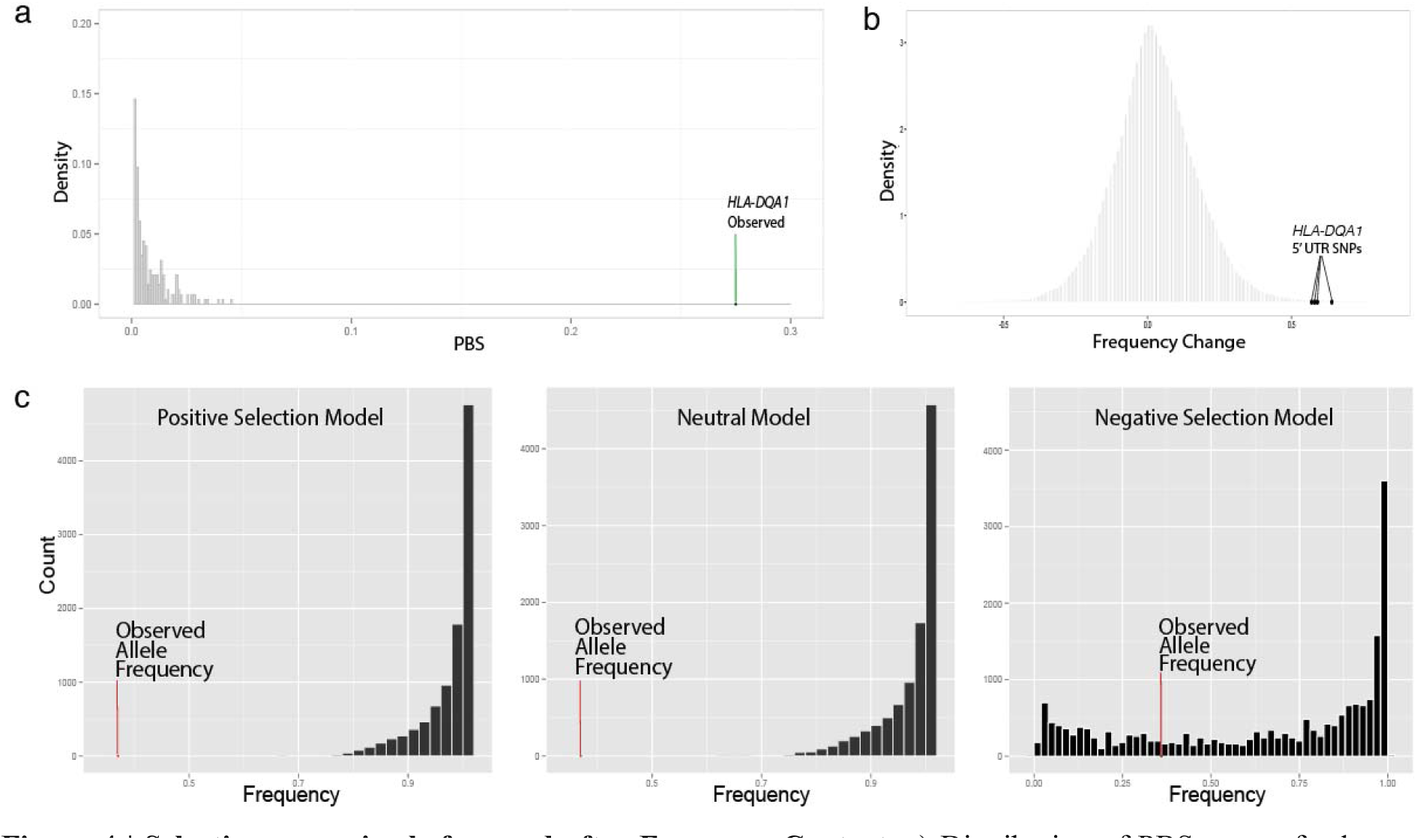
Selection scenarios before and after European Contact. a) Distribution of PBS scores for long-term balancing selection (heterozygote advantage) simulations involving a genomic region of the same length as the *HLA-DQA1* transcript. Given the observed PBS score in the ancient population, this form of balancing selection is inconsistent with our data. (b) Empirical distribution of frequency changes for all derived alleles in the exome between the ancient and modern populations, showing the *HLA-DQA1* UTR5 SNPs as outliers. (c) Simulations of different selection schemes and the allele frequency of the *HLA-DQA1* SNPs observed in the modern Tsimshian. The allele frequency changes between the ancient and modern populations could not be explained by our demographic model (shift due to neutrality after European contact) or one that included only positive selection (no shift from positive selection after European contact). None of the simulations in which the originally positively-selected allele is neutral or still under positive selection after the time of European contact reached the observed allele frequency in the modern population. A model that incorporated negative selection after European contact was a better fit to our data, where ∼26% of the simulations reached the observed frequency.

To model evolutionary forces acting on the *HLA-DQA1* derived alleles after European contact, we chose the frequency of the allele showing the greatest change of 0.67. We utilized a simulation-based approach, described in detail in Supplementary Note 7, to evaluate models under positive, neutral, and negative selection. We also used the same approach to obtain estimates of the correlation between the time of environmental change (*t*) and the selection coefficient (*s*) (Supplementary Fig. 12). Figure 4c shows that neither a neutral or positive selection scheme could fit our data, with none of the simulations reaching the observed frequency in the modern population. However, the negative selection model was compatible, where 26% of the simulations either reached or surpassed the observed frequency.

We also investigated if the observed allele frequency in the ancient population could be better explained by drift rather than selection. Using the same general method, we simulated the initial allele frequencies at the time Native Americans split off from East Asian populations by randomly sampling allele frequencies from a backward simulation conditioned on the modern CHB frequency. We then simulated allele frequencies at 60 generations ago—the time at which the ancient population was sampled. The resulting distribution in Supplementary Figure 7 shows that a neutral scenario is not a good fit for our data.

## Discussion

Our unique data set has allowed us to examine the demography of a single Native American population through three distinct time frames. We first examined the population from a time span of 5,000 years leading to European contact. Selection scans on the ancient individuals from this period revealed a top candidate for positive selection, *HLA-DQA1*, giving the inference of an immune related adaptive event. We next inferred the severity of the population collapse after European contact, which correlates with historical population declines associated with regional smallpox epidemics^44,45^, as well as general estimates of Native American population declines based on mitochondrial DNA diversity^27,46^. During the contact period, previous long standing positive selection on the *HLA-DQA1* gene may also have been significant. The HLA-DQ receptor has been associated with a variety of colonization era infectious disease, including measles^47,48^, tuberculosis^49,50^, and with the adaptive immune response to the vaccinia virus, which is an attenuated form of smallpox^51,52^. Further studies are needed to investigate if the ancient alleles putatively under positive selection may pose a differential disease outcome with respect to European-borne pathogens, as well as their effect on downstream target genes.

However, when examining the population post-contact and into contemporary times, variants of the *HLA-DQA1* gene experience a marked frequency change. This change presents a more complex scenario when taking into account all three time frames. First, scans for positive selection in the modern Tsimshian, with and without correcting for European admixture, revealed no statistically significant selection on immune-related genes (Supplementary Tables 6 and 7). The gene ontology enrichment analyses also did not suggest a correlation with immune function (Supplementary Table 9). Second, demography alone was unable to explain the large frequency change in the *HLA-DQA1* alleles between the ancient and modern groups based on simulations (Fig. 4c). European admixture in the modern individuals also did not account for the frequency changes since the haplotypes in this region can be attributed to Native American ancestry (Supplementary Fig. 15). Furthermore, *HLA-DQA1* remained a top PBS hit in scans involving both a European admixture correction (Supplementary Table 4) and with an additional scan involving unadmixed Native American individuals from a different modern population (suggesting a regional adaptive event) (Supplementary Table 8; ranked fourth best candidate, with the top three functionally uncharacterized).

We therefore explored alternative explanations for the observed frequency change of the *HLA-DQA1* alleles in the time after contact. Since HLA genes have been previously postulated to be under balancing selection in humans^53,54^, we examined the possibility that long-term balancing selection could explain our data by simulating under a model of heterozygote advantage conditional on our inferred demographic model. We found that this specific type of balancing selection is a poor fit to the data, whereby the *HLA-DQA1* gene is still in an extreme outlier relative to the simulation results (Fig. 4a). Next, we used a forward simulation based approach to trace the *HLA-DQA1* allele trajectories under different selection models after the point of European contact. We found that simulations under our demographic model, which was modified to not include European admixture given our local ancestry results (Supplementary Fig. 15b), was insufficient to explain the frequency change in the modern population—with none of the 10^4^ simulations reaching the observed frequency (Fig. 4c). However, upon applying a model of negative selection at the time of contact, we found that simulated allele frequencies were compatible with the observed frequencies in the modern population (Fig. 4c). Although we were unable to precisely identify the selection coefficient necessary to drive the allele frequency change (since the likelihood surface is relatively flat, Supplementary Fig. 12), it is likely that relatively strong negative selection occurred. Such strength would be expected under a time frame of less than 7 generations and correlates with the high mortality rates associated with the regional smallpox epidemics of the 1800s, which reached upwards of 70%^44,45^.

The results presented here reveal an evolutionary history that spans thousands of years. The immune-related alleles that exhibit strong signals of positive selection in the ancient Native Americans from the Northwest Coast, likely correlate to an adaptation to pathogens that were present in the ancient environments of the region. Our results also suggest that these indigenous peoples may have experienced negative selection on the same immune-related genetic component after European contact and the ensuing population collapse. The shift may represent a form of balancing selection due to fluctuating environments^55^. This inference was only made possible through our examination of a single population through time, revealing nuanced demographic events and the utility of such studies. Furthermore, the evolutionary history detailed here helps to better understand the experiences of Native Americans with disease, in both ancient and colonial periods, by demonstrating a shift in immune-related selection pressures associated with the environmental impact of European contact.

## Methods

### Ethics approval and Community Engagement

RSM and JSC visited the partnering First Nations prior to collecting samples for this study. During and after community visits and extensive consultation, a research protocol and informed consent documents—agreed upon by the indigenous communities and the non-indigenous researchers—was approved by the University of Illinois Institutional Review Board (#10538). RSM, JSC and JL visited the community annually during the study to report the latest results and continue to visit the First Nations to report on this and related studies.

### DNA Extraction and Library Preparation

We prepared DNA extracts from 25 ancient individuals from the Prince Rupert region of British Columbia (Supplementary Table 1) and prepared DNA sequencing libraries in a clean room facility. The 25 modern DNA samples underwent similar procedures in a separate facility designated for modern DNA only (Supplementary Note 2).

### Exome Capture

We captured 4 libraries from each ancient sample utilizing the Illumina TruSeq exome enrichment kit, with protocol modifications suited for ancient DNA (*SI Appendix*). The 4 captured libraries from each individual were pooled and sequenced (single-end) on one lane of the Illumina HiSeq 2000. Two libraries from each modern individual were enriched with either the Illumina TruSeq or Nextera exome enrichment kits (Supplementary Table 2) and pooled with an additional modern sample for single-end sequencing on the Illumina HiSeq 2000.

### Contamination Estimates

To estimate contamination across the genome-wide data, we used the ContEst tool^16^. The tool uses a Bayesian approach to calculate both the posterior and the maximum a posteriori probability of contamination level within a BAM file of an individual. This method has been shown effective in detecting contamination in exomes with low coverage^16^. HapMap_3.3 global population frequencies for each SNP, mapped to b37, were used for the estimates. All ancient samples demonstrated contamination below 1%, except for PRH Ancient 163. The estimates are shown on Supplementary Table 3.

### Variant Discovery

Reads below a read length of 35 were filtered out before mapping to hg19. For analyses requiring genotype calls (e.g., *TreeMix, ADMIXTURE)*, SAMtools-1.1^56^ was utilized with a minimum mapping quality of 30, a minimum base quality of 20, a minimum read depth of 6, and a max read depth of 80. Sites were also filtered for violation of a one-tailed test for Hardy-Weinberg Equilibrium at a *p*-value < 10-^4^ ^57^. Due to the low average read depth of the PRH Ancients, genotypes were not called directly for the selection scan. Instead, the program suite ANGSD^58^ was used to compute genotype likelihoods using the SAMtools model and called SNPs via its estimation of allele frequencies. Each alignment used in the estimation was filtered for a minimum mapping quality of 30, a minimum base quality of 20, trimmed at each end for 5 bp to minimize biases from DNA deamination, and a minimum p-value threshold of 10^−6^.

### PBS Selection Scan

To detect regions under positive selection in both the PRH Ancient and the Tsimshian, the population branch statistics (PBS)^33^ was utilized. The population branch statistics (PBS) has proven powerful in detecting hypoxia adaptation in high altitude populations ^33,34^. It employs a set of three populations (call them *X, Y*, and *Z*), and assumes that they have the rooted relationship ((*X,Y*),*Z*). In actuality, the calculation for the PBS does not require a rooted tree, and so their specific rooted relationship does not matter.

An analogous statistic can be calculated for populations *Y* and *Z*. In this study, we are concerned with the situation in which *X* is the PRH Ancients, *Y* is the modern Tsimshian, and Z is the Han Chinese (CHB from the 1,000 Genomes Project). We therefore are interested in computing:

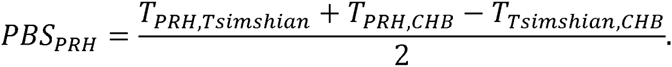

Because the *ADMIXTURE and TreeMix* analyses (Figs 1a and 1c) indicate a likely admixture event between the modern Tsimshian and Europeans, the allele frequencies in the modern Tsimshian were corrected for admixture using the method described by Huerta-Sánchez et al. ^34^. Let 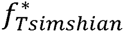 and ƒ_*Tsimshian*_ represent the allele frequency at a locus in the Tsimshian population pre and post admixture, respectively. Further, assuming that we use Europeans as a proxy, we let ƒ_*European*_ be the allele frequency at the same locus in a reference European population (we used the Great Britain GBR population from the 1,000 Genomes Project). Assuming that the proportion of ancestry derived from Europeans at the locus is *α*, under a model of instantaneous admixture, the allele frequency in the Tsimshian post admixture would be

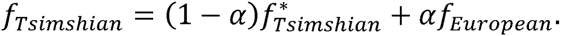

Rearranging, we can solve for the allele frequency prior to admixture as

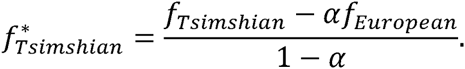

We estimated *α* at each locus by choosing an *α* value that minimized the *F*_*ST*_ between the admixture-corrected Tsimshian and the Han Chinese (CHB) outgroup population. We used the admixture-corrected allele frequencies for the scans for positive selection.

We used *ANGSD* ^58^ to compute allele frequencies for the modern Tsimshian, the PRH Ancient, the Han Chinese (CHB), and the Great Britain (GBR) populations directly from the raw sequencing reads, accounting for the uncertainty in genotype calling. Allele frequencies were based on 25 individuals from each population. We required that reads had a map quality of at least 30 and each nucleotide had a quality of at least 20. We also only called allele frequencies at sites in which data from at least five individuals was not completely missing. To additionally guard against post-mortem deamination, we trimmed the first and last five nucleotides of each read in the PRH Ancient samples. The allele frequencies in the modern Tsimshian were subsequently corrected for potential European ancestry (see procedure in the directly preceding paragraph).

We performed two scans for positive selection. The first was a per-gene scan, in which we calculated PBS for the PRH Ancient population (*PBS*_*PRH*_) using all data at given gene. Gene annotations were derived from RefSeq, utilizing the longest transcript for a given gene. The transcript length was taken as the transcription start to the transcription stop, and included both introns and exons. *F*_*ST*_ between each pair of populations was calculated using all SNPs that fell between the transcription start and stop of the gene, as well as 10 kilobases upstream of the transcription start and 10 kilobases downstream of the transcription stop (similar to how it was performed in Huerta-Sánchez et al.^34^). We then ranked each gene in the genome with decreasing PBS. The top two candiates are displayed in Table 2.

The second scan was a per-SNP scan, in which we calculated PBS for the PRH Ancient population (*PBS*_*PRH*_) and the modern Tsimshian population (*PBS*_*Tsimshian*_) at each SNP. That is, *F*_*S*T_ between each pair of populations was calculated for a given SNP, and this set of *F*_*ST*_ values was used to calculate PBS for that SNP. We then created Manhattan plots, and highlight chromosome 6 in Figure 3.

We also performed an analogous scan in which we substituted 25 mostly unadmixed Peruvian samples from the 1,000 Genomes Project Phase 3^41^ for the 25 modern Tsimshian individuals. We identified the individuals showing little to no admixture by running *ADMIXTURE* ^25^. The individuals identified and used for this and subsequent analyses were: HG01572, HG01923, HG01926, HG01927, HG01941, HG01951, HG01953, HG01954, HG02008, HG02102, HG02105, HG02146, HG02147, HG02150, HG02259, HG02260, HG02266, HG02271, HG02272, HG02275, HG02278, HG02291, HG02292, HG02304, HG02348. Because the individuals appeared mostly unadmixed, we did not correct allele frequencies for admixture. Results from this scan are highlighted in Supplementary Table 10. Using his different reference sister Native American population still suggests that *HLA-DQA1* is a reasonable candidate, as it is ranked fourth in the scan—with the three genes ranked above it devoid of functional characterization—using Peruvians rather than modern Tsimshian as a reference sister population.

### Demographic History Model

Parameters for the demographic model (Fig. 2) were inferred with FastSimCoal2^32^. The fixed parameters were implemented from Gravel et al. and were as follows: out of Africa bottleneck (*N* =1861, *T*=51kya) ^31^, split between the CHB and GBR (serving as the ghost population) (*N*_*GBR*_=1032, *N*_*CHB*_=550; *T*=23kya) ^31^. Admixture between the GBR and Tsimshian (*T*=100 years, admixture fraction=0.33) were taken from Verdu et al. ^24^. One hundred optimizations were run for the inferred values, taking the best likelihood parameters from each of the 100 sets. The data was simulated with an effective sequence length of 7.4 Mb and per-base per-generation mutation and recombination rates of 2.5×10^−8^. The optimizations utilized joint derived site frequency spectra (SFS) for the CHB, PRH Ancients, and Tsimshian. The European population (Great Britain denoted by GBR) served as a ghost population in the model. This SFS contained 7.4 Mb of monomorphic and polymorphic sites based on hg19 potential synonymous sites, where data was reported for each individual. A parametric bootstrapping approach was used to construct the 95% confidence intervals. The inferred parameters and confidence intervals are listed in Table 1 and the model residuals are depicted in Supplementary Figure 13.

## Acknowledgments

This project was made possible through the active collaboration of the Lax Kw’alaams and Metlakatla First Nations. We would also like to thank Jun Li for furnishing comparative exome data from contemporary Native American populations. The research was funded by the National Science Foundation (#BCS-1413551) and by the Office of the Vice Chancellor of Research, University of Illinois at Urbana-Champaign, by the Canadian Museum of History, Gatineau, Quebec, Canada, and by Pennsylvania State University startup funds. Portions of this research were conducted with the Advanced CyberInfrastructure computational resources provided by The Institute for CyberScience at Pennsylvania State University.

### Author Contributions

Conceived and designed the study RSM, JL, MD. Performed the experiments: JL and MR. Analyzed the data: JL, MD, EHS, SN. Contributed reagents/materials/analysis tools: RSM, JL, MD, EW. Wrote the paper: JL, RSM, MD, JSC with contributions from all authors. Community engagement: RSM, JSC, BP, JM, JL. Discussed and interpreted results: JL, MD, RSM, EHS.

### Additional information

**Accession codes:** The ancient data have NCBI Short Read Archive accession no. 288803. The data from modern individuals are available via a data access agreement with RSM at the University of Illinois.

**Supplementary Information** is linked to the online version of the paper at www.nature.com/naturecommunications.

